# Hallucination of moving objects revealed by a dynamic noise background

**DOI:** 10.1101/2020.08.21.262170

**Authors:** Ryohei Nakayama, Alex O. Holcombe

**Affiliations:** Center for Information and Neural Networks (CiNet), National Institute of Information and Communications Technology, 1-4 Yamadaoka, Suita City, Osaka, 565-0871, Japan; School of Psychology, The University of Sydney, NSW 2006, Australia

## Abstract

We show that on a dynamic noise background, the perceived disappearance location of a moving object is shifted in the direction of motion. This “twinkle goes” illusion has little dependence on the luminance- or chromaticity-based confusability of the object with the background, or on the amount of background motion energy in the same direction as the object motion. This suggests that the illusion is enabled by the dynamic noise masking the offset transients that otherwise accompany an object’s disappearance. While these results are consistent with an anticipatory process that pre-activates positions ahead of the object’s current position, additional findings suggest an alternative account: a continuation of attentional tracking after the object disappears. First, the shift was greatly reduced when attention was divided between two moving objects. Second, the illusion was associated with a prolonging of the perceived duration of the object, by an amount that matched the extent of extrapolation inferred from the effect of speed on the size of the illusion (~50 ms). While the anticipatory extrapolation theory does not predict this, the continuation of attentional tracking theory does. Specifically, we propose that in the absence of offset transients, attentional tracking keeps moving for several tens of milliseconds after the target disappearance, and this causes one to hallucinate a moving object at the position of attention.

## Introduction

Objects are not always perceived at the location corresponding to the position they stimulate on the retinas. Object motion, for example, can strongly influence perceived object location (motion-induced position shift or MIPS; De Valois & De Valois, 1991; Ramachandran & Anstis, 1990). A popular theory of such phenomena has been that they reflect anticipatory extrapolation by the brain to compensate for neural delays (Nijhawan, 1994). Other explanations are viable, however. For example, at least some motion and position illusions may be better explained by adaptive integration of uncertain position and motion signals (Fu, Shen, & Dan, 2001; Kwon, Tadin, & Knill, 2015).

Anticipatory extrapolation predicts that if an object suddenly disappears, its disappearance location should be perceived ahead of where it actually disappeared. But this does not occur (Kerzel, 2000; Kerzel, Jordan, & Müsseler, 2001; Whitney & Cavanagh, 2002; Whitney, Murakami, & Cavanagh, 2000). To explain this, advocates of the anticipatory extrapolation theory have suggested that the abrupt disappearance and associated luminance transient results in a correction or reset of extrapolation (Blom, Feuerriegel, Johnson, Bode, & Hogendoorn, 2020; Hogendoorn, 2020; Nijhawan, 2002, 2008). We will refer to this as the *transient correction* hypothesis. A major concern about the transient correction hypothesis is that it appears to have been created post hoc to preserve the extrapolation theory. However, Nijhawan & colleagues (Maus & Nijhawan, 2008; Shi & Nijhawan, 2012) found novel support for the transient-correction hypothesis by assessing the perceived location of objects after they entered the retinal blindspot. The perceived disappearance location of objects moving into the blindspot was beyond the blindspot’s proximal edge. This is consistent with the transient correction theory, because entering the blindspot prevents the disappearance of the moving object from causing a luminance transient. Nijhawan et al. concluded that the perceived disappearance location of an object moving into the blindspot reflects anticipatory activation of positions (in this case, inside the blindspot) ahead of the object’s currently sensed location. That is, throughout the time that the object was presented, it was perceived ahead of its sensed location, and its movement into the blindspot prevented the transient correction process that normally occurs upon object disappearance.

Here we find support for an alternative theory consistent with Nijhawan’s results and that also explains a striking new position and motion illusion that we call the “twinkle goes” illusion. Our theory is that attentional tracking can continue after a moving object’s disappearance and result in continuing perception of the object as attention continues to move beyond the location of the object’s disappearance.

In our demonstration of the new twinkle goes illusion, two smoothly moving objects are presented on a static background for one interval, and on a dynamic noise background during a second interval (Supplementary Movie S1). In the dynamic background portion of the movie, when the moving objects disappear, they appear to be shifted in the direction of motion, whereas on the static background portion they do not.

The results of our first experiments show that the twinkle goes illusion occurs even when the moving objects and background are distinguished with different colors (Experiment 1) and with high luminance contrast (Experiment 2), suggesting the illusion is not due to simple confusion of the background with the moving object. The results also show that for the illusion to occur, the background must be dynamic during a short interval after the moving object’s disappearance. The illusion does not depend on the presence of motion signals in the dynamic background compatible with the motion of the object (Experiment 3). These findings indicate that the offset transient that normally accompanies an object’s disappearance prevents the illusion in normal viewing – the dynamic noise enables the illusion by masking the offset transient.

The results of Experiment 4 show that the size of the illusion increases approximately linearly with speed, up to over 30 deg/s. The approximate linearity is consistent with a process that shifts the perceived location of the object to where it would be in a certain amount of time, specifically 56 ms (*SEM* = 25 ms) after disappearance. Experiment 5 indicates that the twinkle goes illusion is highly dependent on the availability of attention, suggesting that attentional tracking (Pylyshyn & Storm, 1988) may be critical for the phenomenon.

To explain these findings, we propose a *tracking continuation* theory of the illusion. On this theory, attention continues moving in the direction the target was moving, for dozens of milliseconds after the object disappears. This attentional tracking process provides a top-down prediction that causes hallucination of the moving object in the associated positions. Without a dynamic background, this does not occur because attention is captured by the luminance transient associated with an object’s disappearance (Hollingworth, Simons, & Franconeri, 2010; Kawahara, Yanase, & Kitazaki, 2012).

A critical prediction of tracking continuation theory is that the moment that the object is perceived to disappear should be after its actual disappearance. More specifically, to correspond with the amount of time attention must continue tracking to explain the 50 to 60 ms position shift, the object should be judged to disappear 50 to 60 ms after its actual disappearance. Our final experiment (Experiment 6) therefore assessed perceived disappearance time with temporal order judgements. The participants judged the time of disappearance relative to a sound that occurred at a variable time. The results indicate that the perceived disappearance time with a dynamic background is 50 ms (*SEM* = 13 ms) later than that with a static background.

In summary, the twinkle goes illusion appears to be caused by the continuation of attentional tracking in the absence of a salient offset transient when an object disappears. Remarkably, this implies that the attentional tracking process can cause one to hallucinate an object in novel locations.

## Results

### Experiment 1

The perceptual extrapolation apparent in the twinkle goes illusion might conceivably happen because the human visual system confuses luminance components contained in the dynamic noise with a part of the moving objects, resulting in an apparent extension of their trajectory. We tested whether the illusion occurred when the moving objects had a color not shared by the background, and found that the illusion still occurred (Figure S1) – the magnitude of the shift was positive (compared to zero, *z* = 2.53, *p* = 0.006) with the dynamic background and greater than that on the static background (*t*_(5)_ = 5.39, *p* = 0.003, *d_z_* = 2.20), where it was not statistically significant (*z* = −0.47, *p* =0.68) and may have been zero or negative (positive numbers indicate a shift in the direction of object motion). This result indicates that the illusion does not depend on simple confusability of the objects with their background.

### Experiments 2a and 2b

To better understand the role of the dynamic noise background, these experiments varied the time when the dynamic noise was presented relative to the time of the disappearance of the object. In Experiment 2a, the onset time of the background manipulation was varied for each of two different target luminances and in Experiment 2b, the offset time was varied as well as the onset time.

The amount of perceived shift for the disappearance location is plotted in Figure 2, as a function of timing between disappearance and when the background became dynamic rather than static (Experiment 2a). The result shows that when the dynamic noise precedes or coincides with the object disappearance and continues until the end of each trial, the illusion occurs (*z*s > 2.07, *p*s < 0.04), but does not occur (*z*s < 0.83, *p*s > 0.41) when the dynamic noise does not start until 80 ms after the disappearance or is totally absent (static background).

**Figure 1.**
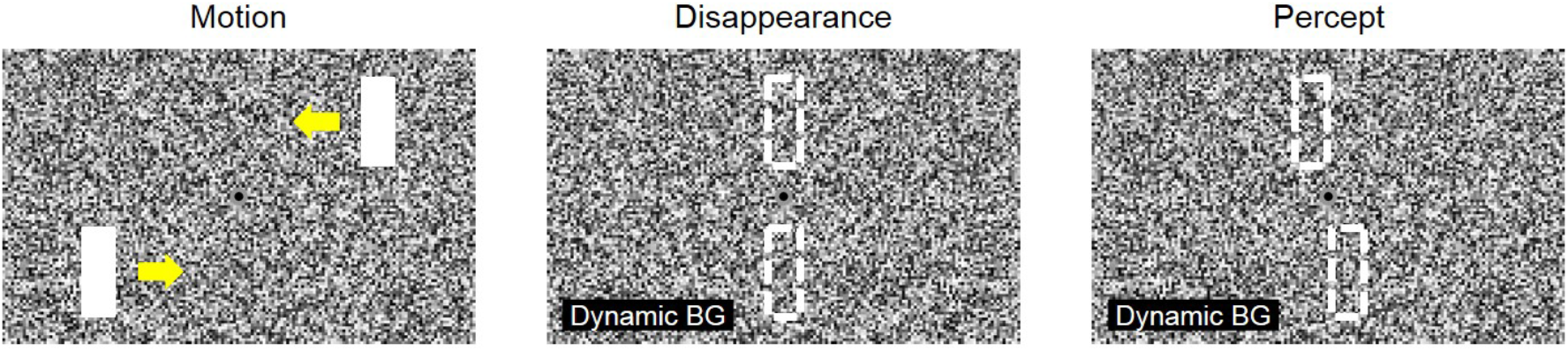
Schematic display of stimuli used here to elicit the “twinkle goes” illusion. One object in the upper visual field and another in the lower visual field moves horizontally and abruptly disappears near the display’s center. Yellow arrows (not shown in the actual display) indicate the movement directions, and dashed rectangles indicate the physical disappearance location (middle panel) and perceived position (right panel), which is shifted in the direction of motion if the random-dot background is dynamically modulated in luminance after the object disappearance. In the experiments, the magnitude of the shift was estimated with alignment judgements where a staircase (1 up 1 down) adjusted the misalignment across trials.

**Figure 2.**
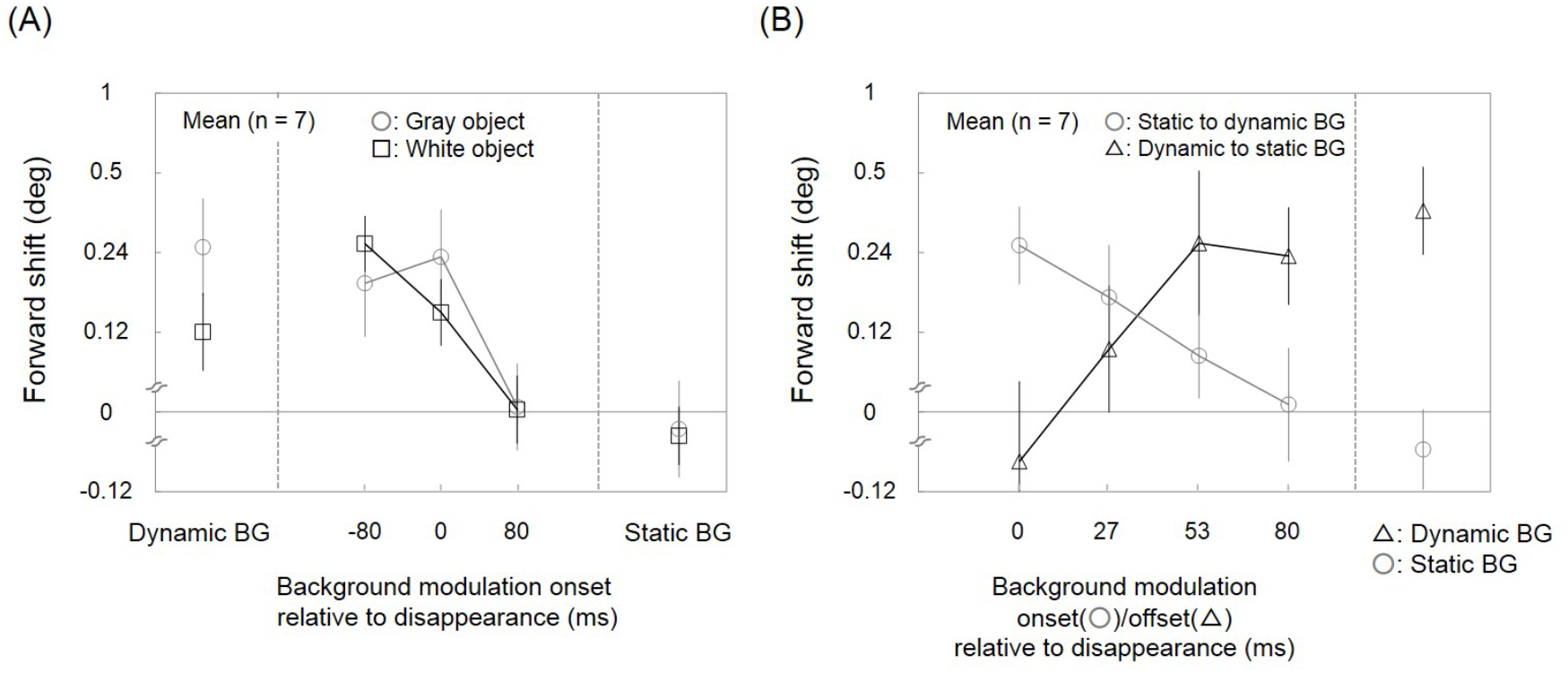
Results of Experiments 2a and 2b. The abscissa shows when the background underwent luminance modulation. The ordinate shows the average shift in perceived position of the object disappearance in the motion direction. The label of dynamic or static BG indicates the presence or absence of the background modulation throughout each trial. Other numbers in the abscissa indicate when the background modulation started (displayed by circles or triangles) or stopped (displayed by triangles; tested only in Experiment 2b) relative to the object disappearance. The shift for gray objects (low luminance contrast with the background) is displayed by circles or triangles and the shift for white objects (high luminance contrast) is displayed by squares (tested only in Experiment 2a). Error bars represent ± *SEM*.

Although the different target luminance values (white and gray) had substantially different saliences with the background, this had little effect on the illusion, further supporting the conclusion of Experiment 1 that the object need not be highly confusable with the background (Figure 2A; main effect of object luminance, *F*_(1, 6)_ = 1.12, *p* = 0.33, η_p_^2^ = 0.01, in a two-way repeated measures ANOVA). The illusion was somewhat smaller when the objects were white (interaction of object luminance with background being static vs. dynamic, *F*_(4, 24)_ = 7.55, *p* = 0.0004, η_p_^2^ = 0.04; simple main effect of luminance when the background was dynamic throughout: *F*_(1, 30)_ = 10.46, *p* = 0.003, η_p_^2^ = 0.03). This may stem from the disappearance of the objects against the dynamic background being more salient when the objects are white than when they are gray. That is, the ability of the dynamic noise to mask the transient and thus cause the illusion is reduced when the object is white, although the illusion was still present: the shift was larger on the dynamic background than on the static background (*t*_(48)_ = 7.38, *p* < 0.0001, *d_z_* = 2.79 for gray objects; *t*_(48)_ = 4.22, *p* = 0.0001, *d_z_* = 1.60 for white objects; main effect of background being static vs. dynamic, *F*_(4, 24)_ = 27.20, *p* < 0.0001, η_p_^2^ = 0.33).

Experiment 2a revealed that the critical period for the background being dynamic to create the illusion did not extend more than 80 ms after the time of the object disappearance. Experiment 2b extended these results by also investigating the effect of the duration of the dynamic aspect of the background, by varying both its onset and offset time.

In half of trials, the background was dynamic at the end of the trial, but the time when that dynamic modulation started was varied, more finely than in Experiment 2a. In the other half of trials, the background was dynamic at the beginning of the trial, but when it ceased was varied. The results revealed (Figure 2B, circles) that the shift decreases as the dynamic noise starts later (one-way repeated measures ANOVA, *F*_(4, 24)_ = 11.73, *p* < 0.0001, η_p_^2^ = 0.66), falling to approximately zero by 80 ms after the objects’ disappearance (*z* = 0.13, *p* = 0.90). This indicates that the background must be dynamic for at least some time after the objects’ disappearance, as it seems that no shift occurs if ongoing dynamic noise ceases at the same time as the disappearance (*z* = 0.62, *p* = 0.54; Figure 2B, triangles). As the time the background is dynamic after the object disappearance is extended, the shift increases, for several dozen milliseconds (*F*_(4, 24)_ = 12.99, *p* < 0.0001, η_p_^2^ = 0.68).

In summary, the illusion is strongest when the background is dynamic for the 80 ms following object disappearance. The more that the background is dynamic during that interval, the greater the perceptual shift.

### Experiment 3

In the conditions studied so far, the dynamic background contained an equal amount of motion components, or energy, in all directions. Conceivably, the motion energy in the object movement direction might drive the effect, as the expectation of motion in the forward direction might cause attention to weight that stimulus component more heavily. To assess this possibility, the magnitude of the extrapolation was estimated with objects moving on a dynamic background filled with orthogonal movement of the dots rather than random luminance modulation.

The illusion still occurred (Figure S2). A positive shift was found (*z* = 1.91, *p* = 0.03) with the moving background but not the static background (*z* = 0.10, *p* = 0.46), producing a significant effect of the dynamic/static background (*t*_(5)_ = 3.12, *p* = 0.02, *d_z_* = 1.27).

Rather than the motion energy of the dynamic background or confusability with the moving objects being critical to its effect, there must be some other mechanism. We speculate that the dynamic noise enables the illusion by masking the luminance transient associated with the object disappearance.

### Experiments 4a and 4b

Experiments 4a and 4b investigated the effect of speed on the amount of perceptual shift. Horizontally moving objects (Experiment 4a, Figure 3) were tested for speeds up to 36.2 deg/s. The shift increased with speed (one-way repeated measures ANOVA, *F*_(2, 16)_ = 46.70, *p* < 0.0001, η_p_^2^ = 0.52) and the effect of speed was approximately linear – the slope from 2.3 deg/sec to 9.0 deg/sec, 0.89 (*SEM* = 0.19), was similar to the slope from 9.0 deg/sec to 36.2 deg/sec, 0.80 (*SEM =* 0.10), and not statistically significantly different, *t*_(8)_ = 0.16, *p* = 0.88, *d_z_* = 0.29. The overall slope implies 56 ms (*SEM* = 25 ms) of extrapolation.

**Figure 3.**
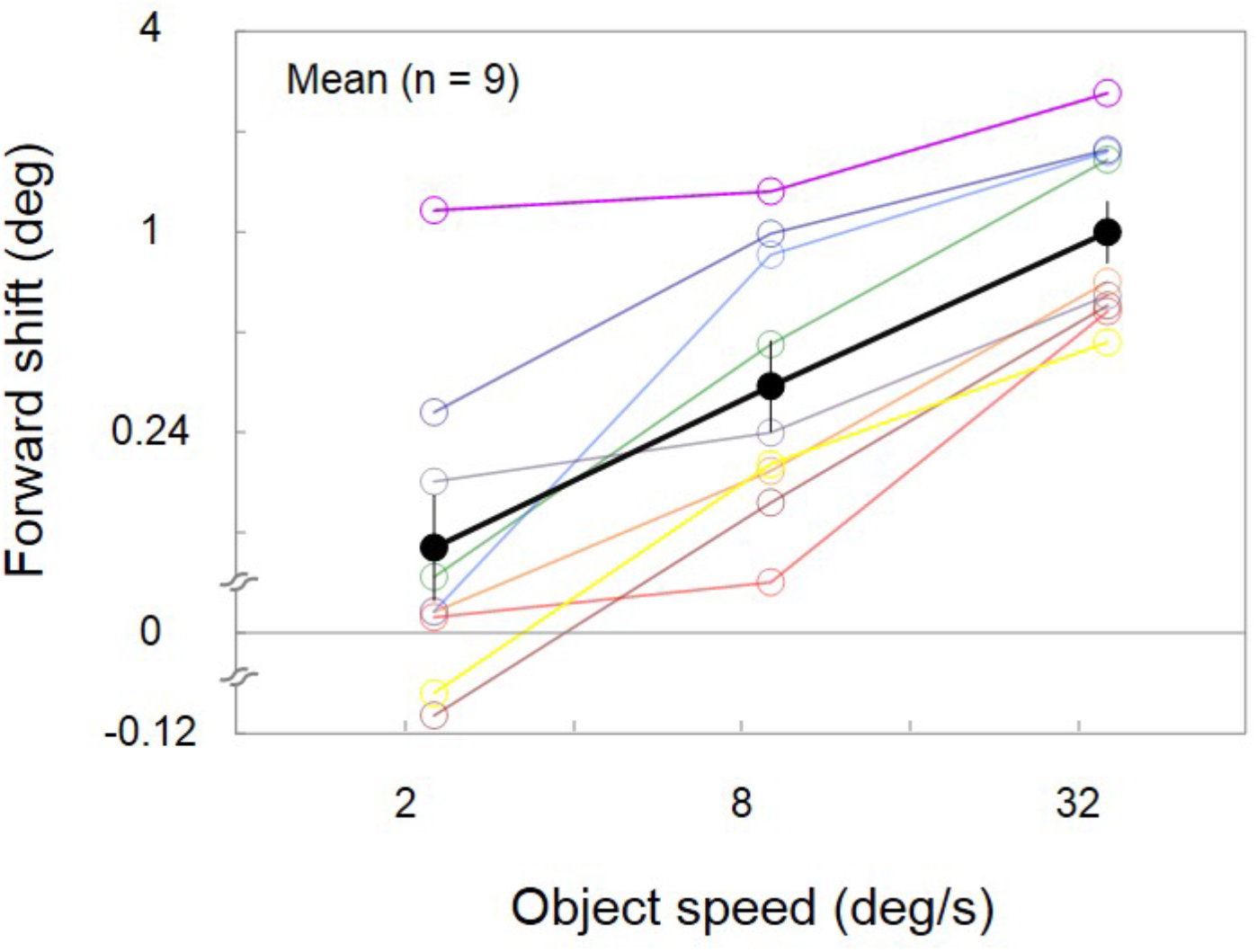
Results of Experiments 4a. The perceived forward shift from the disappearance location is shown as a function of object speed in horizontal motion. Empty and filled circles display individual and average results respectively. Error bars represent ± *SEM*.

The effect of speed is quite distinct from speed’s effect on the MIPS phenomenon, which saturates at low speeds (Bressler & Whitney, 2006). The approximate linearity of the twinkle goes over a wide range of speed is consistent with a process that extrapolates the objects’ position by a further 50-60 ms after object disappearance. To assess the role of attentional tracking, we tested objects that rotated about fixation (Experiment 4b), because previous work has found that participants can only attentionally track such objects up to about 2 revolutions per second, with performance declining for some people already at 1.6 rps (Holcombe & Chen, 2012; Verstraten, Cavanagh, & Labianca, 2000). Here we found that the illusory position shift increased with speed (one-way repeated measures ANOVA up to 1.2 rps, *F*_(3, 18)_ = 6.45, *p* = 0.004, η_p_^2^ = 0.26; Figure 4) until about 1.6 rps (corresponding to 87.2 deg/s), consistent with the possibility that the ability to attentionally track constrains the effect. Note that unlike the previous Experiment (4a), a particular shift amount only has meaning when compared to that of other speeds, because it is the amount of shift tangential to the objects’ circular trajectory, as it is derived from participants’ judgements of vertical misalignment.

**Figure 4.**
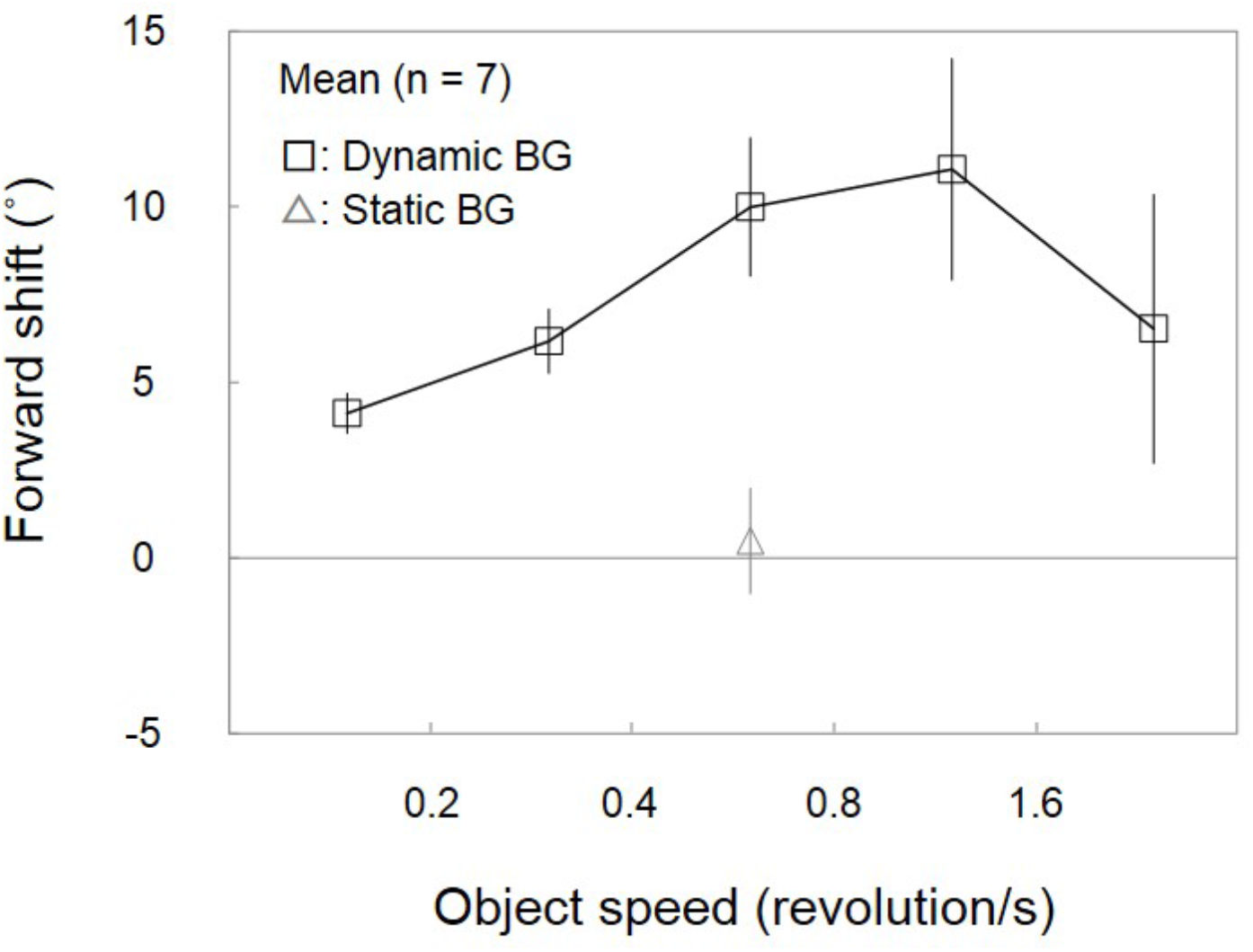
Results of Experiments 4b. The perceived forward shift from the disappearance location is shown as a function of object speed in revolution. The background underwent luminance modulation in five-sixths of the trials (dynamic BG; displayed by squares), with no modulation in the other trials (static BG; displayed by a triangle). Error bars represent ± *SEM*. One revolution/s corresponds to 54.5 deg/s.

### Experiment 5

The result of Experiment 4b suggested that attentional tracking may play a critical role in the extrapolation of a moving target. This would be consistent with work on multiple object tracking, which has found evidence that a consistent object trajectory benefits tracking performance. This extrapolation process that underlies multiple object tracking is resource-limited; the more targets tracked, the less the benefit of a consistent motion trajectory (Howe & Holcombe, 2012; Luu & Howe, 2015).

Here we investigated whether the degree of extrapolation in the twinkle goes illusion also depends critically on the amount of attention resource devoted to a moving target. To manipulate whether undivided attention was on the moving targets, we added a second pair of objects to the display (Figure 5A) and varied whether participants knew which pair they would have to judge the final position of at the end of the trial.

**Figure 5.**
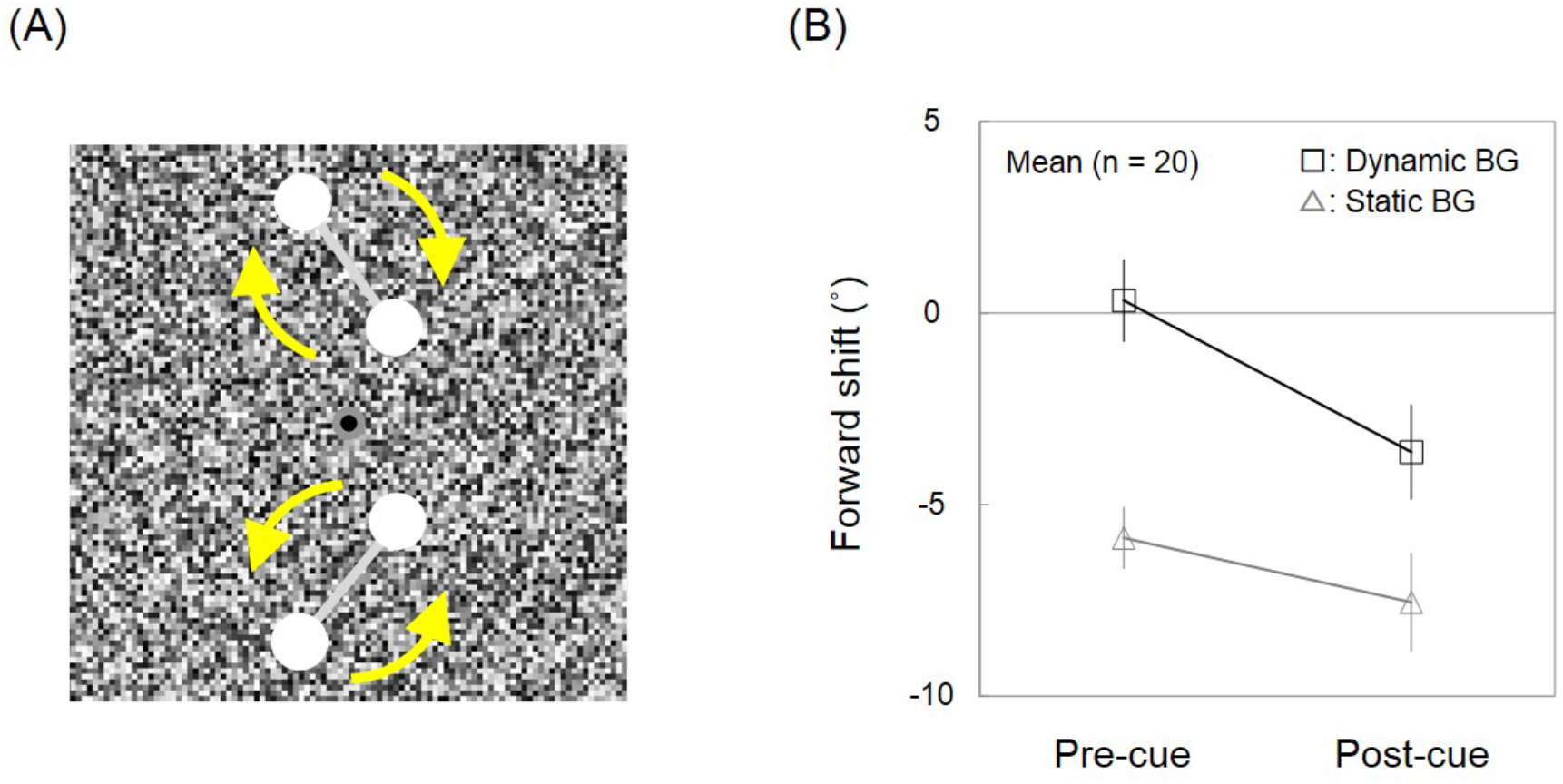
Schematic stimulus display and results of Experiment 5. (A) Two pairs of circular objects revolve clockwise and counter-clockwise as indicated by yellow arrows (not shown in the actual display) and then disappear. The target pair was indicated both before and after the stimulus presentation in half of trials (pre-cue condition), and only after the stimulus presentation in the other half of trials (post-cue condition). (B) The forward shift in the disappearance location is shown for the pre-cue and the post-cue conditions. The background underwent luminance modulation in half of trials (dynamic BG; displayed by squares) and did not in the other half of trials (static BG; displayed by triangles). Error bars represent ± *SEM*.

In half of trials, the target pair was indicated before the stimulus presentation (the pre-cue condition), so participants could attend solely to that pair. In the other half of trials, the target pair was not indicated until after (the post-cue condition), so participants must split their attention to track both pairs, which impairs tracking performance (Alvarez & Cavanagh, 2005; Holcombe & Chen, 2013; Pylyshyn & Storm, 1988). Rather than splitting their attention to track both pairs, each with half the resource, they might attend to what turns out to be the target pair in half of trials, and the non-target pair in the other half of trials, but previous work has found that at speeds like that used here, participants typically attempt to track both (Chen, Howe, & Holcombe, 2013).

The results reveal the strong effect of availability of attention, supporting a critical role for attentional tracking. In the post-cue condition, where only half the attention resource was available, the perceived position of the target with the dynamic background significantly lagged its final position (negative shift, *z* = −2.93, *p* = 0.004; Figure 5), while not in the pre-cue condition (*z* = 0.31, *p* = 0.76). Two-way repeated measures ANOVA confirms the basic main effect of the pre-/post-cue (*F*_(1, 19)_ = 11.42, *p* = 0.003, η_p_^2^ = 0.08) and that of the dynamic/static background (*F*_(1, 19)_ = 52.76, *p* < 0.0001, η_p_^2^ = 0.21). A simple main effect analysis supported the effect of pre/post cue in the dynamic background condition (*F*_(1,38)_ = 18.14, *p* = 0.0001, η_p_^2^ = 0.08). With the static background, the pre-/post-cue had less effect, as indicated by the significant interaction with dynamic versus static background, *F*_(1, 19)_ = 7.73, *p* = 0.01, η_p_^2^ = 0.01; the simple main effect of pre/post cue in the static background condition was not statistically significant, *F*_(1,38)_ = 3.26, *p* = 0.08, η_p_^2^ = 0.01.

Significant lags like that found in the post-cue conditions, and on the static background, are thought to reflect intermittent sampling of position by attentional tracking (Holcombe & Chen, 2013; Howard & Holcombe, 2008; VanRullen, Carlson, & Cavanagh, 2007). On our attentional tracking continuation account, at the time of disappearance the last sampled position may be substantially behind the actual disappearance position, but in the dynamic background condition, attentional tracking can continue for several dozen milliseconds, reducing or eliminating the lag.

### Experiment 6

On the tracking continuation theory, the perceived disappearance location is shifted because attentional tracking continues to move after the target disappearance. This theory predicts that the disappearance of the moving object will be perceived to occur 50-60 ms later than in conditions where the illusion does not occur. This is unlike the anticipatory extrapolation theory, where the purpose of the shift is to reduce or eliminate the lag caused by neural delay, so the shifted perceived position is not something created after object disappearance.

To test the contrasting predictions of the anticipatory theory and the tracking continuation theory, the moment that the disappearance was perceived to occur was estimated using an acoustic probe (a click sound). The task was to judge whether the click was presented before or after the disappearance.

The results revealed that the perceived disappearance time of the moving object was later for the dynamic backgrounds, where the illusion occurred, than for the static (*t*_(6)_ = 3.78, *p* = 0.009, *d_z_* = 1.43; Figure S3). The dynamic background did not hinder participants’ ability to discriminate temporal order (i.e., slope of the fitted logistic curve), *t*_(6)_ = 1.96, *p* = 0.10, *d_z_* = 0.74, consistent with a Cass & Van der Burg (2014) result that nearby dynamic noise did not impair temporal order judgements between auditory and visual stimuli. The perceived disappearance time for every participant was later for the dynamic background than for the static background, on average by 50 ms (*SEM* = 13 ms), which corresponds approximately to the amount of extrapolation found in Experiment 4a. This result supports the tracking continuation rather than the anticipatory extrapolation hypothesis. One way the masking of the offset transient may result in the effect may be to delay the detection of the disappearance of the moving object, causing tracking to continue until the object disappearance is detected.

## Discussion

The twinkle goes illusion reveals that with a dynamic background, the perceived disappearance location of a moving object is shifted forward. The illusion does not require the moving object to be highly confusable with the background, nor does it depend on a large amount of motion energy in the background in common with the object movement direction.

We propose that the dynamic noise enables the illusion by masking the offset transient associated with the disappearance of the object. In this we agree with Nijhawan that offset transients prevent perception of a moving object in an extrapolated position (Nijhawan, 2002, 2008). However, Nijhawan’s theory was an anticipatory prediction theory, that moving objects are continually represented in an extrapolated position ahead of the corresponding sensory signal, and it is that anticipatory representation that is perceived in the absence of an offset transient. Other theorists have also suggested that illusory perceptual shifts in the direction of object motion are caused by an anticipatory extrapolated representation. That is, neurons corresponding to locations ahead of a moving object are activated in advance, giving rise to perception of the moving object in that location before it reaches that location (Blom et al., 2020).

Neural activation at anticipated locations certainly does occur (e.g., Berry, Brivanlou, Jordan, & Meister, 1999), and under certain circumstances, such as high spatial uncertainty, such activation may be made perceptually manifest so that the object is perceived ahead of its currently-sensed location (Fu, Shen, Gao, & Dan, 2004; Whitney et al., 2003). For the twinkle goes illusion, however, it appears that the percept occurs after the cessation of afferent sensory signals from the moving object, and that it reflects a continuation of attentional tracking along the object trajectory. One study showed that visual sensitivity is enhanced in front of the leading edge of a drifting grating pattern (Roach, McGraw, & Johnston, 2011), which we suggest could be due to the continuation of attentional tracking, although in that case we know of no evidence against the anticipatory neural activation account.

Because low-level motion mechanisms show speed or temporal frequency tuning rather than a monotonic effect of speed (Burr & Ross, 1982; Kelly, 1979; Perrone & Thiele, 2002), the linear increase in shift we observed suggests the amount of shift is determined by a high-level motion mechanism, namely attentional tracking. Further supporting this theory is that the shift declines when object speed nears the tracking limit, and that perceived positions increasingly show lag rather than extrapolation when attention is divided. Finally, the time the object is perceived to disappear is later than its physical disappearance time, by an amount (~50 ms) consistent with the amount of extrapolation. We propose that if not captured by the offset transient, attentional tracking can continue for 50-60 ms on average, causing one to hallucinate a moving object in the corresponding positions.

### Relation to visual persistence

An alternative account is that participants might observe visual persistence for 50-60 ms after the moving object disappears. However, any persisting images would not be expected to move when participants maintain fixation (Kerzel, 2000; Kerzel et al., 2001). Moreover, visual persistence should be suppressed immediately by a surface (here, the dynamic background) filled with transient luminance changes (Castet, 1994; Francis, 1996). Although we did not record participants’ eye movements, linear eye movements could not explain the illusion because the objects judged move in opposite directions. While torsion could occur, it would be difficult to explain why eye movements (e.g., torsion) would differ with static and dynamic backgrounds.

### Relation to anticipatory extrapolation

Previous researchers have suggested that phenomena such as the flash-lag effect are caused by anticipatory extrapolation – activation of neurons ahead of an object’s currently sensed position (e.g., Hogendoorn, 2020; Nijhawan, 2008). But both the flash-lag effect and other illusions involving a shift of perception in the direction of motion have been shown to be determined largely by motion after the perceptual probe rather than before (Brenner & Smeets, 2000; Cavanagh & Anstis, 2013; Eagleman & Sejnowski, 2000; Roach & McGraw, 2009) This supports postdictive explanations or asynchronous binding explanations (Cai & Schlag, 2001) rather than one based on perception of neurons pre-activated ahead of a moving object.

Nijhawan and colleagues addressed the postdictive property of the flash-lag effect by suggesting that transients or motion signals, when they occur after the flash, prevent perception of a pre-existing extrapolated representation. Howe et al. (2008) already undermined this account by showing that adding transients to the moving object (both before and after the flash) do not eliminate the flash-lag effect. Nijhawan’s account is further undermined by our documenting here an extrapolated percept exposed by masking of offset transients that seems to arise after the disappearance of the afferent signals, not before.

### Relation to adaptive integration of uncertain position and motion signals

Kwon et al. (2015) suggested a computational model of MIPS based on Kalman filtering that accounts well for various quantitative characteristics of MIPS. Their model calculates object position by integrating sensory position signals with current motion signals attributed to the object, weighted by position uncertainty relative to motion uncertainty. Kwon et al. conceptualize attentional tracking as the integration locus of low-level motion signals with position signals. However, their model would not expect a significant extrapolation for our stimuli because our stimuli have no pattern motion that could be attributed to position change. On the other hand, perhaps the model takes some time to stop tracking when a moving object disappears. This lag might correspond to a continuation of attentional tracking.

On the framework of adaptive integration of uncertain signals, however, a linear increase in shift with speed would not be expected (e.g., Fu et al., 2001; Kanai, Sheth, & Shimojo, 2004) partly because the visual system has a slow speed prior (Stocker & Simoncelli, 2006; Weiss, Simoncelli, & Adelson, 2002) that underestimates object speed (and position shift as a result) particularly for high speeds, especially because fast objects should yield more position uncertainty (Brenner, Van Beers, Rotman, & Smeets, 2006; Linares, Holcombe, & White, 2009).

The present illusion has different characteristics from the MIPS phenomenon that the uncertainty-based theory has been successful in explaining. While dividing attention reduced the twinkle goes, previous work showed that for the MIPS, dividing attention and even the absence of awareness of the motion has little effect (Haladjian, Lisi, & Cavanagh, 2018; Nakayama & Holcombe, 2020; Whitney, 2005). In addition, Kwon et al. (2015) found that when a moving object is shifted in position by orthogonal pattern motion, the abrupt disappearance of the object on a static background still results in the shift being perceived, rather than eliminated by the offset transient. The visual system may bias position estimates to compensate for positional uncertainty, which may chiefly cause the MIPS, but the present results support a distinct mechanism for the twinkle goes illusion

### Role of attentional tracking

Top-down processes, including attentional tracking, have been shown in other contexts to contribute to position coding (Shim & Cavanagh, 2005; Whitney, 2006). The localization of a flash is repelled away from the location of pre-cuing (Suzuki & Cavanagh, 1997), while it is attracted toward the location of post-cuing (Ono & Watanabe, 2011), suggesting a role for attentional shifts. This is in agreement with neurophysiological findings of remapping of neuronal receptive fields around the locus of attention (Connor, Preddie, Gallant, & Van Essen, 1997; Womelsdorf, Anton-Erxleben, Pieper, & Treue, 2006). A related mechanism to distort perceptual space may contribute to the present illusion, but does not suffice because our temporal order judgment experiment indicated that the moving object is extrapolated not only in space but also in time.

Attention enhances sensory processing in cortex (Brefczynski & DeYoe, 1999; Liu, Pestilli, & Carrasco, 2005; Martínez-Trujillo & Treue, 2002; Reynolds, Pasternak, & Desimone, 2000). One study reported illusory percepts elicited when the location of a visual stimulus is cued hundreds milliseconds after the stimulus disappeared (Sergent et al., 2013), suggesting attention can re-evoke bottom-up processes for conscious perception. Top-down prediction is nevertheless critical for the present effect because it occurs at a location that has been never stimulated by the object. Consistent with this notion, a recent study decoding EEG found that attentional tracking shares in part multivariate patterns of neural signals with perceptual mechanism (Robinson et al., 2020). The dynamic noise required for the twinkle goes illusion might provide some supporting sensory signals for visibility (reminiscent of the phonemic restoration effect; Warren, 1970), although this possibility is somewhat undermined because the illusion little depends on the resemblance of object and background stimuli.

## Conclusion

The present study indicates that top-down prediction due to attentional tracking can in this instance cause a percept. The object’s disappearance may provide a prediction error signal (Rao & Ballard, 1999) that eliminates the continuation of prediction.

While predictive coding and related approaches have been popular, we are not aware of other psychophysical evidence for an attention-created highly suprathreshold percept. However, other literature has found evidence that imagery can bias perception to create faint percepts (Waller, Schweitzer, Brunton, & Knudson, 2012), voluntary attention can group stimuli into new objects (Ongchoco & Scholl, 2019), and unusual circumstances such as sensory restriction can result in hallucinations that may reflect top-down predictions (Miskovic et al., 2019).

## Methods

### Participants

Six observers (1 female, 1 author) participated in both Experiment 1 and Experiment 3. Seven observers (6 female, 1 author) participated in Experiment 2a and a different seven observers (5 female) participated in Experiment 2b. Nine observers (7 female, 1 author) participated in Experiment 4a and seven observers (3 female, 1 author) participated in Experiment 4b. Twenty observers (8 female, 1 author) participated in Experiment 5. Seven observers (1 female, 1 author) participated in Experiment 6. All participants had normal or corrected-to-normal vision and provided written informed consent. All experiments were conducted in accordance with the Declaration of Helsinki (2003) and were approved by the ethics committee of the University of Sydney (Australia) or the National Institute of Information and Communications Technology (Japan).

### Apparatus

Images were displayed on a gamma-corrected 22-inch CRT screen (1280×1024 pixel) with a frame rate of 75 Hz. The CRT resolution was 3.6 min/pixel at a viewing distance of 30 cm.

### Experiment 1

A pair of rectangular objects (2.9 deg wide x 7.7 deg high) moved horizontally at 18.1 deg/s from opposite sides toward the center, in upper and lower visual fields, with a vertical distance of 7.7 deg from a black fixation point (0.5 deg in diameter), respectively (as described in Figure 1). The central area of the background (77.1 deg wide x 36.2 deg high), except a circular gray part (1.1 deg in diameter) around the fixation point, was filled with square dots (0.2 x 0.2 deg).

The moving objects and the background were completely defined by blue and red components. Either color was randomly assigned to the objects and the other color to the background, i.e., if the background was blue, only the blue gun of the CRT screen was used for the background, with the red and green set to zero (black), and only the red gun was used for the objects. The luminance of the object was 15.0 cd/m^2^ for red (0.62, 0.35 in CIE 1931 xy chromaticity) and 9.9 cd/m^2^ for blue (0.15, 0.08). The luminance values of the background square dots were randomly chosen from a uniform distribution between 0 and 9.9 cd/m^2^ for blue and between 0 and 15.0 cd/m^2^ for red and, in dynamic background conditions, refreshed every movie frame (75 Hz).

The moving objects were presented for a random duration between 0.8-1 s and then disappeared at pre-determined positions, after which the background was presented for an additional 400 ms. The background luminance modulation (which did not occur in the static background condition) started 80 ms before the object disappearance (and continued 400 ms after the disappearance).

In each trial, participants viewed the stimulus display with steady fixation and reported whether the direction of vertical misalignment at disappearance of moving objects was to the left (top object offset to the left of the bottom object) or to the right. A blank (uniform gray) display was presented during the response period and after that a new static noise background with a fixation point was presented for 800 ms until the next trial started. The physical misalignment was adjusted by a staircase with a factor of 2 (e.g., −1.8, −0.9, −0.5, −0.2, 0, 0.2, 0.5, 0.9, 1.8 deg) and a 1 up 1 down rule, which targets a 50% proportion of “right” direction responses, with 120 trials per condition.

The dynamic and static background conditions were randomly interleaved along with corresponding staircases. The motion directions of the upper and lower visual fields were swapped every trial. For each condition, we estimated the point of subjective equality (PSE) as the vertical alignment corresponding to chance reporting of the direction, by fitting a logistic curve via maximum likelihood.

### Experiments 2a & 2b

The same stimuli as Experiment 1 were used except that the moving objects and the background were achromatic. The luminance values of the background square dots were randomly chosen from a uniform distribution between 0.3 and 113.5 cd/m^2^. In Experiment 2a, the background was static or dynamic throughout the stimulus presentation, or the modulation started 80 ms before, at the same time with (0 ms after), or 80 ms after the object disappearance. The objects were either white (high luminance contrast with the background) or gray (low luminance contrast with the background). In Experiment 2b, while the objects were invariably gray, the background was static or dynamic throughout, or the modulation started 0, 27, 54, or 80 ms after the disappearance, or the modulation started at the same time with the motion onset then stopped 0, 27, 54, or 80 ms after the disappearance (background changed from dynamic to static).

These conditions were randomly interleaved along with corresponding staircases (1200 trials in each experiment). Otherwise, measurements and analyses were done in the same way as Experiment 1.

### Experiment 3

The same stimuli as Experiment 2 were used except that the dynamic background was a moving background: instead of refreshing luminance values, half of the square dots filling the background area moved upward and the other half moved downward, at 9.0 deg/s. The dots that had reached either top or bottom edge of the background area (67.5% of dots did not reach the edge during the stimulus presentation on average) disappeared and new dots appeared at the other edge. The background motion (which did not occur in the static background condition) started 80 ms before the disappearance, and continued until 400 ms after the disappearance. Measurements and analyses were done in the same way as Experiment 1.

### Experiments 4a and 4b

Experiment 4a used the same horizontal motion of gray rectangle objects as Experiment 2. The object speed was 2.3, 9.0, or 36.2 deg/s and the background was dynamic throughout the stimulus presentation.

Experiment 4b used a pair of white circular objects (2.4 deg in diameter) that revolved about the fixation point with a radius of 8.7 deg and a speed of 0.15, 0.3, 0.6, 1.2, or 2.4 revolution/s. The direction of revolution for each trial was the opposite of the previous trial. The background modulation started 80 ms before the disappearance, and continued until 400 ms after the objects’ disappearance. A static background condition was also included, with only 0.6 revolution/s tested. Other aspects were same as for Experiment 2 except that the objects were presented for a random time between 0.8-1.6 s and the background area subtended 48.2 deg x 48.2 deg.

Measurements and analyses were done in the same way as Experiment 1, except in Experiment 4b the physical misalignment of revolving objects was adjusted by a staircase with a stepsize of 4° in polar angle (e.g., −16°, −12°, −8°, −4°, 0°, 4°, 8°, 12°, 16°) and a 1 up 1 down rule with 120 trials per condition.

### Experiment 5

Two pairs of white circular objects (2.4 deg in diameter) revolved clockwise and counter-clockwise at 1.3 revolution/s in upper and lower visual fields, with a radius of 4.3 deg, respectively (Figure 5A). The centroids of the object pairs were 13.5 deg above and below the fixation point. As a reference for alignment judgments, each pair was connected by a gray thin line (0.5 deg wide and 6.3 deg high).

Half of trials were the pre-cue condition and half were the post-cue condition, which appeared in random order. At the beginning of a pre-cue trial, a white vertical line (0.5 deg wide and 3.9 deg high) was shown for 1.2 s either 2.9 deg above or below the fixation point to cue the location where the target pair would appear (upper or lower visual field). In the post-cue condition trials, two cuing lines were shown both above and below to indicate that the participant needed to attend to both pairs. After the stimulus presentation, in all trials the white vertical line indicating the pair to report the alignment of was shown until the response. Both conditions were further divided into dynamic versus static background conditions, which were randomly interleaved, with the final misalignment of each controlled by its own staircase (480 trials in total). The revolving directions were swapped every trial and the target pair was assigned randomly. The physical misalignment of the distracter pair was determined at random between −16° and 16°. Measurements and analyses were otherwise same as Experiment 4b.

### Experiment 6

A gray circular object revolved about the central fixation at 0.6 revolution/s. The initial (and final) position was randomized and for each trial, the revolving direction was different from the previous trial. The background was static or dynamic throughout the stimulus presentation. Other aspects were same as for Experiment 4b. As an acoustic probe stimulus, a 5-ms click was presented via a headphone at different times relative to the time of object disappearance.

In each trial, participants viewed the stimulus display with steady fixation and reported whether the click was presented before or after the disappearance of the moving object (2AFC). A blank (uniform gray) display was presented during the response period and after that a new static noise background with a fixation point was presented for 800 ms until the next trial started. The time of the presentation of the click relative to the disappearance of the moving object was adjusted by a staircase with a step of 53 ms (4 movie frames) and a 1 up 1 down rule, which targets a 50% proportion of “click first” responses, with 120 trials per condition. For each condition, we estimated the point of subjective simultaneity (PSS) by fitting a logistic curve via maximum likelihood.

## Acknowledgments

We thank Chaz Firestone for discussions. Supported by JSPS KAKENHI Grant JP18J01398 to RN. This study was carried out when RN was a JSPS research fellow; a visiting researcher at CiNet, NICT; and a visiting researcher at the University of Sydney.

## Author Contributions

RN and AOH designed the study. RN performed experiments and analyzed data. RN and AOH wrote the manuscript.

## Competing interests

The authors declare no competing interests.

## Supplementary Figures and Movie

**Figure S1.**
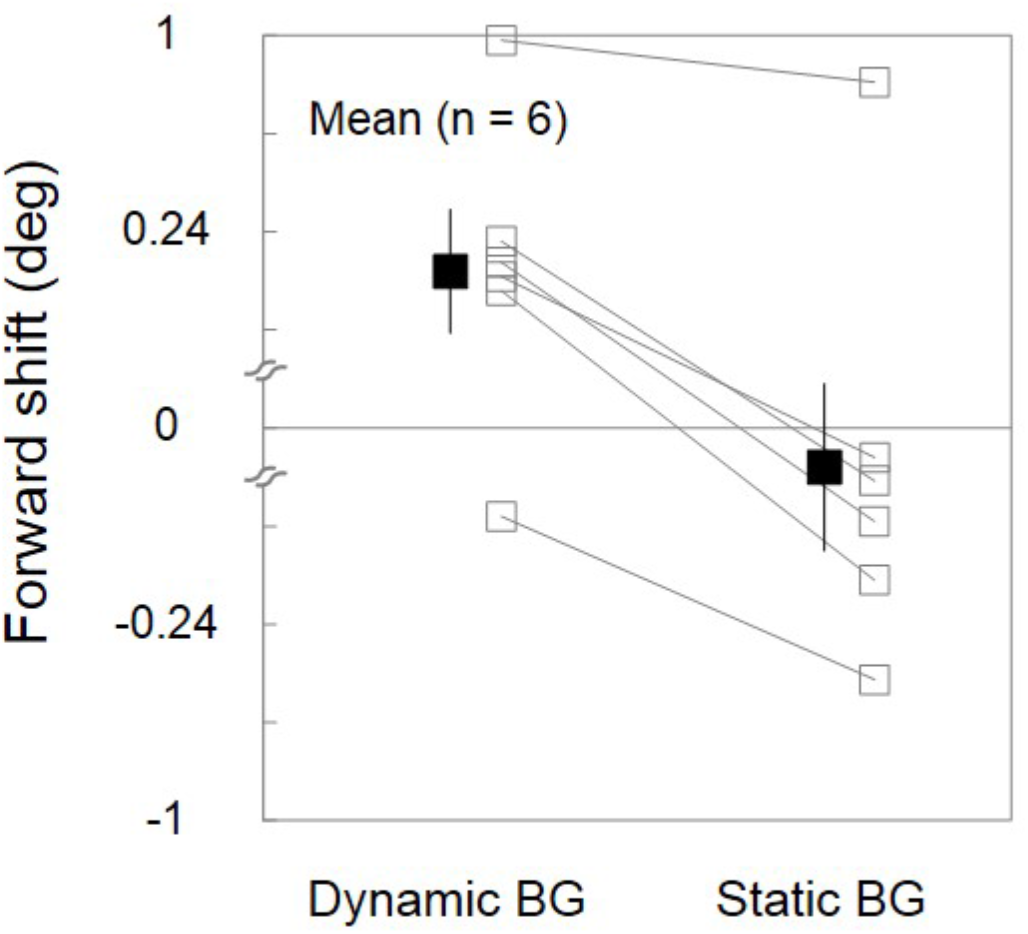
Results of Experiment 1. The objects had a different color from their background. The forward shift in the disappearance location is shown for each of the dynamic and static backgrounds. Empty and filled squares display individual and average results respectively. Error bars represent ± *SEM*.

**Figure S2.**
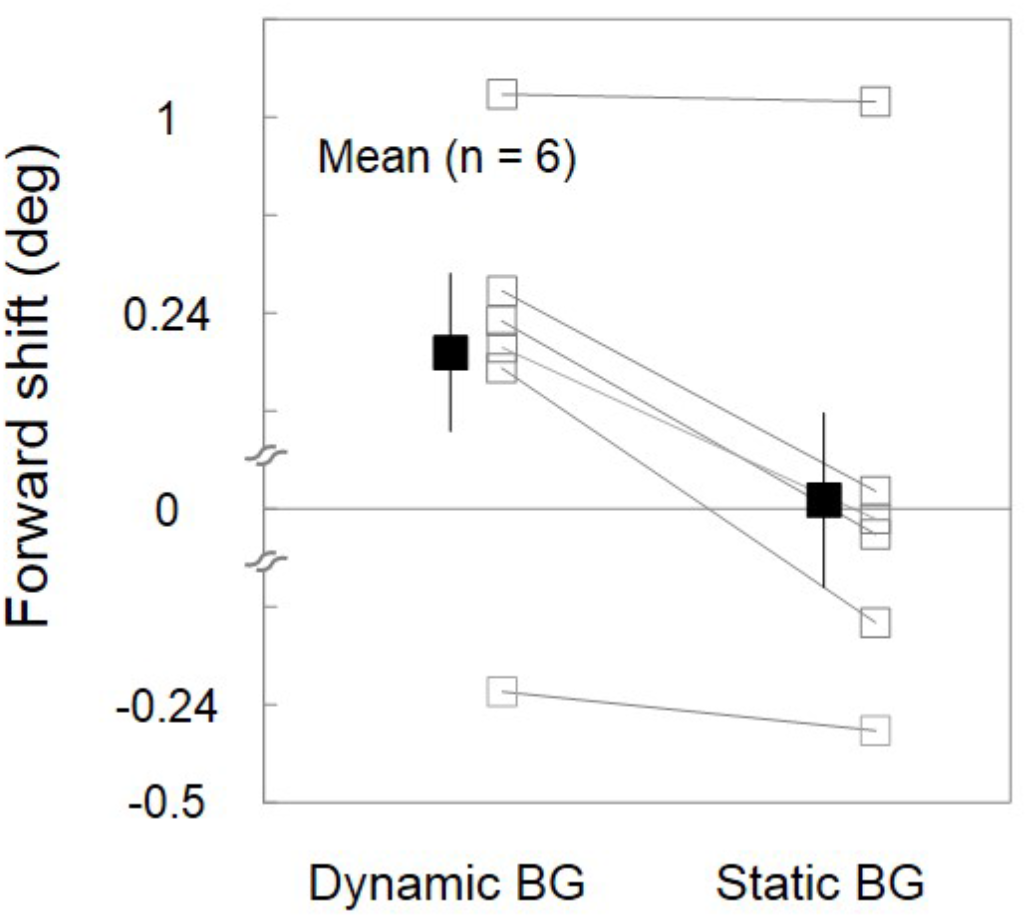
Results of Experiment 3. The dynamic background was entirely composed of vertically moving dots whereas the objects moved horizontally. The forward shift in the disappearance location is shown for each of the dynamic and static backgrounds. Empty and filled squares display individual and average results respectively. Error bars represent ± *SEM*.

**Figure S3.**
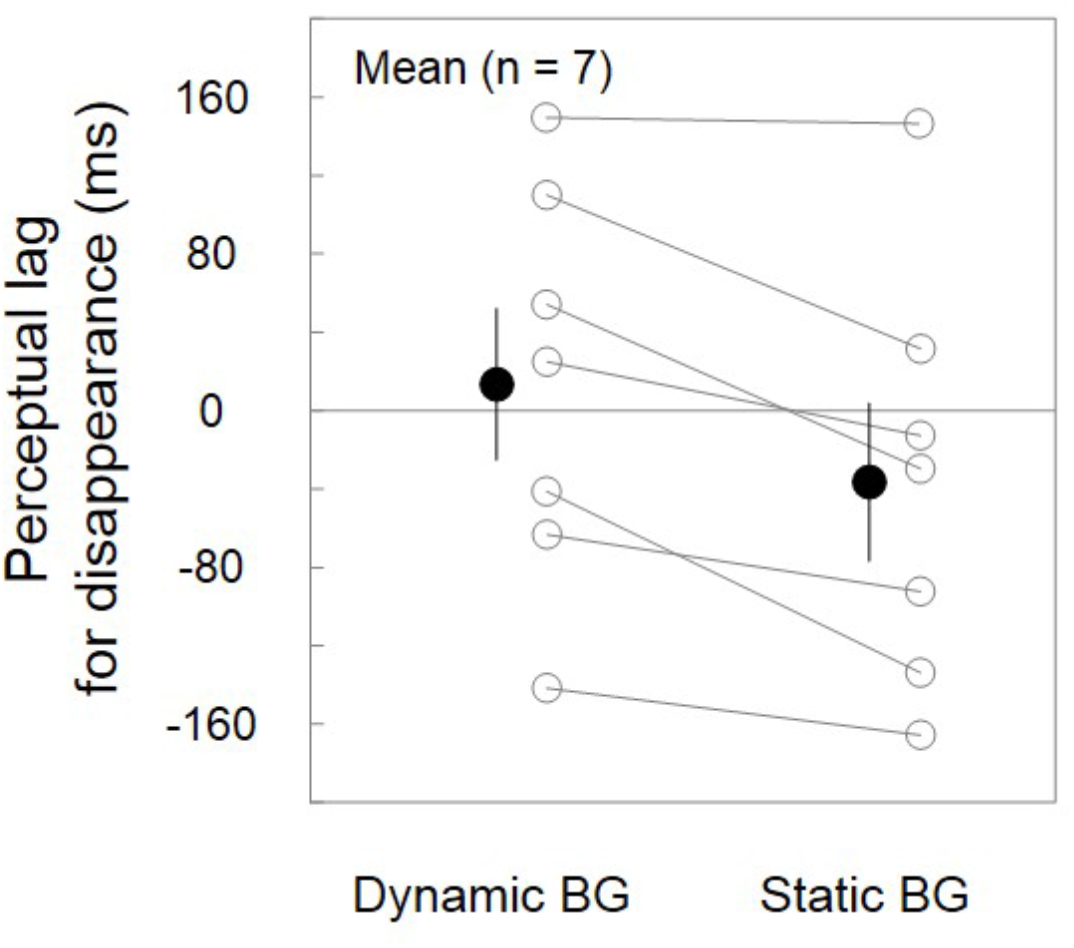
Results of Experiment 6. The subjective time for the disappearance of a moving object, estimated by temporal order judgements with an acoustic probe, relative to the actual disappearance time is shown for each of the dynamic and static backgrounds. Empty and filled circles display individual and average results respectively. Error bars represent ± *SEM*.

**Movie S1.**
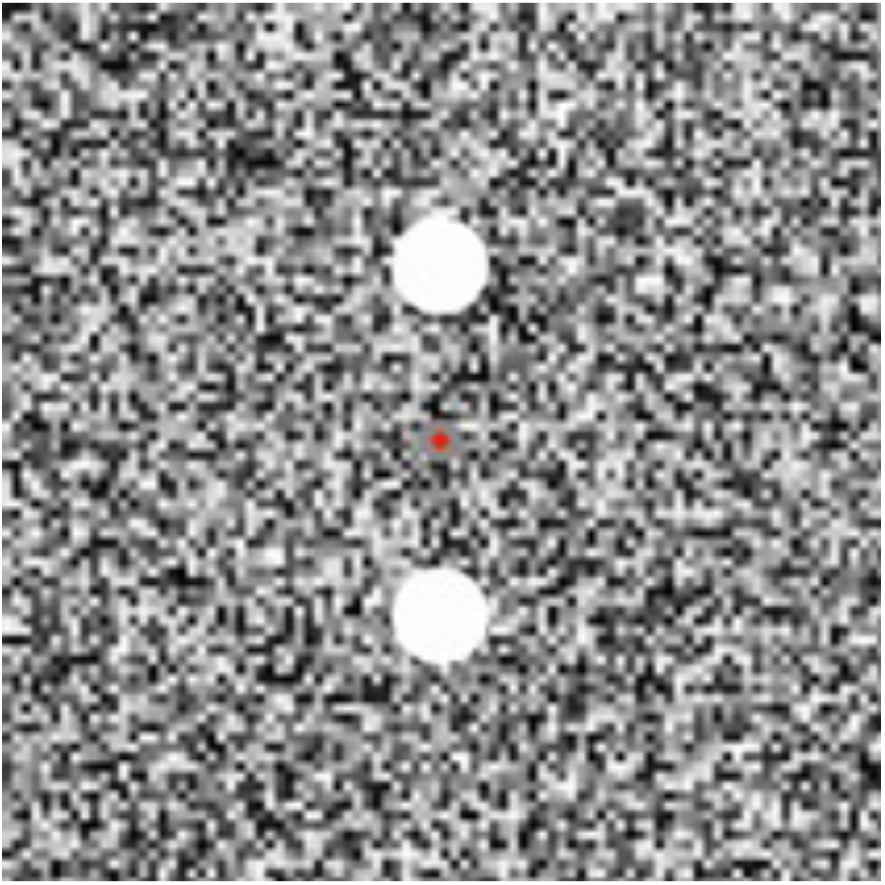
“Twinkle goes”, a new motion illusion of extrapolation. The revolving discs disappear when they are vertically aligned. But in the part of the movie where the disappearance is followed immediately by twinkle (dynamic noise), the discs appear shifted in the direction of motion. This does not occur when the background remains static.

## Notes

### Competing Interest Statement

The authors have declared no competing interest.

